# Genome-wide Marginal Epistatic Association Mapping in Case-Control Studies

**DOI:** 10.1101/374983

**Authors:** Lorin Crawford, Xiang Zhou

## Abstract

Epistasis, commonly defined as the interaction between genetic loci, is an important contributor to the genetic architecture underlying many complex traits and common diseases. Most existing epistatic mapping methods in genome-wide association studies explicitly search over all pairwise or higher-order interactions. However, due to the potentially large search space and the resulting multiple testing burden, these conventional approaches often suffer from heavy computational cost and low statistical power. A recently proposed attractive alternative for mapping epistasis focuses instead on detecting marginal epistasis, which is defined as the combined pairwise interaction effects between a given variant and all other variants. By searching for marginal epistatic effects, one can identify genetic variants that are involved in epistasis without the need to identify the exact partners with which the variants interact — thus, potentially alleviating much of the statistical and computational burden associated with conventional epistatic mapping procedures. However, previous marginal epistatic mapping methods are based on quantitative trait models. As we will show here, these lack statistical power in case-control studies. Here, we develop a liability threshold mixed model that extends marginal epistatic mapping to case-control studies. Our method properly accounts for case-control ascertainment and the binary nature of case-control data. We refer to this method as the liability threshold marginal epistasis test (LT-MAPIT). With simulations, we illustrate the benefits of LT-MAPIT in terms of providing effective type I error control, and being more powerful than both existing marginal epistatic mapping methods and conventional explicit search-based approaches in case-control data. We finally apply LT-MAPIT to identify both marginal and pairwise epistasis in seven complex diseases from the Wellcome Trust Case Control Consortium (WTCCC) 1 study.

## Introduction

Epistasis, commonly defined as the interaction between genetic loci, has long been thought to play a key role in defining the genetic architecture underlying many complex traits and common diseases [1, 2]. Indeed, previous studies have detected pervasive epistasis in many model organisms [3–26]. Substantial contributions of epistasis to phenotypic variance have been revealed for many complex traits [27, 28] and have been suggested to constitute the genetic basis of evolution [29, 30]. Furthermore, modeling epistasis in addition to additive/dominant effects has been shown to increase phenotypic prediction accuracy in model organisms [31] and facilitate genomic selection in animal breeding programs [32, 33]. Despite some controversies [34], recent genetic mapping studies have also identified candidates of epistatic interactions that significantly contribute to quantitative traits and diseases [35–39]. Importantly, epistasis has been proposed as a key contributor to missing heritability — the proportion of heritability not explained by the top associated variants in GWASs [40–43].

The recent development of genome-wide association studies (GWASs) [42] has provided an unique opportunity to search across the whole genome to detect genetic variants that are involved in epistasis. Many statistical works have been developed to facilitate the identification of epistasis in GWASs [44, 45]. Generally, these existing tools can be classified into two categories. The first category of methods is conventional and explicitly searches for pairwise or higher-order interactions when identifying epistatic effects. These approaches rely on distinct criteria for selecting a testing unit based on variants or genes [46]. In particular, they use different searching strategies including exhaustive search [47–49], probabilistic search [50], or prioritization based on a preselected candidate set [51]. Specific statistical paradigms have been implemented for explicit search-based approaches including various frequentist tests [52] and Bayesian inferences [53–55]. Unfortunately, because of the extremely large search space (e.g. *p*(*p* − 1)/2 possible pairwise combinations for *p*-variants), these methods often suffer from heavy computational burden. Computationally, despite various efficient computational improvements [48, 56] and recently developed search algorithms [50], exploring over a large combinatorial domain remains a daunting task for large epistatic mapping studies. Statistically, because of a lack of *a priori* knowledge about epistatic loci, exploring all combinations of genetic variants could result in low statistical power — on the other hand, restricting to a subset of prioritized combinations based on prior knowledge or marginal additive effects could also miss important genetic interactions.

The second category of epistatic mapping methods, exemplified by the recently developed MArginal ePIstasis Test (MAPIT) [39], attempts to address the previously mentioned challenges by alternatively detecting *marginal* epistasis. Specifically, instead of directly identifying individual pairwise or higher-order interactions, these approaches focus on identifying variants that have a non-zero interaction effect with any other variants. For example, MAPIT tests each variant (in turn) for its *marginal epistatic effect* — the combined pairwise interaction effects between a given variant and all other variants. By testing marginal epistatic effects, one can identify candidate markers that are involved in epistasis without the need to identify the exact partners with which the variants interact — thus, potentially alleviating much of the statistical and computational burden associated with the first category of epistatic mapping methods. In addition, evidence of significant marginal epistasis detected by MAPIT can be used to further prioritize the search for meaningful pairwise interactions. Despite its potential advantages, however, MAPIT has previously been restricted to only quantitative traits. As we will show below, a naïve application of MAPIT to case-control studies yields surprisingly low power. Thus, an overall restriction to quantitative traits limits the potential impact of marginal epistatic mapping methods.

Here, we develop a new statistical method that allows us to detect marginal epistasis in case-control studies. Our method relies on a liability threshold mixed model [57–59] to properly account for case-control ascertainment and the binary nature of case-control data. By combining efficient variance component estimation and hypothesis testing procedures [39,60] together with reasonable model approximations, our proposed approach is scalable to moderately sized GWASs. We refer to this method as the liability threshold marginal epistasis test (LT-MAPIT). With simulations, we illustrate the benefits of LT-MAPIT in terms of providing effective type I error control, as well as being more powerful than both existing marginal epistatic models and conventional search-based approaches in case-control data. Lastly, we use LT-MAPIT to identify both marginal and pairwise epistasis in seven complex diseases from the Wellcome Trust Case Control Consortium (WTCCC) 1 study [61].

## Material and Methods

### Modeling Marginal Epistasis for Case-Control Data

We begin by briefly reviewing the notation and intuition behind the MArginal ePIstasis Test (MAPIT) for case-control studies. Our goal is two-fold. First, we aim to identify variants that interact with other variants without having to explicitly search over the entire model space of all possible interactions. Second, we aim to properly account for case-control ascertainment and the binary nature of case-control data. To achieve these objectives, we consider a liability threshold mixed effects model. Here, we assume that the case-control status *y_i_* for the *i*-th individual is determined by the their corresponding underlying liability value *l_i_*. Namely,

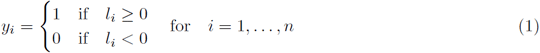

where *y_i_* = 1 if the *i*-th individual is a case, and *y_i_* = 0 when the *i*-th individual is a control. It is important to note that while the disease status **y** = (*y*_1_, …, *y_n_*)^T^ is binary, the underlying liability **l** = (*l*_1_, …, *l_n_*)^T^ is continuous. The liability threshold model allows us to adapt classical statistical methods, that were initially developed for quantitative phenotypes, and use them to analyze the unobserved continuous trait of liability [57–59]. In our case, we examine one variant at a time (indexed by *r*), and consider the following regression

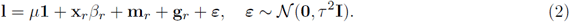

where *μ* is an intercept term multiplied with an *n*-dimensional vector of ones **1**; **x**_*r*_ is an *n*-dimensional genotype vector for the *r*-th variant that is the focus of the model; *β_r_* is the corresponding additive effect size; 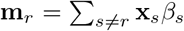 is the combined additive effects from all other *s* ≠ *r* variants with effect sizes *β_s_*, and effectively represents the polygenic background of all variants except for *r*-th 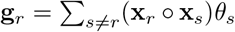 is the summation of all pairwise interaction effects **x**_*r*_ ○ **x**_*s*_ between the *r*-th variant and all other variants *s* ≠ *r* with regression coefficient *θ_s_*; ***ε*** is an *n*-vector of residual errors; *τ* ^2^ is the residual error variance; **I** denotes the identity matrix; and ***N*** (•,•) denotes a multivariate normal distribution. The term **g**_*r*_ is the main focus of the model and represents the marginal epistatic effect of *r*-th variant — formally defined as the summation of its epistatic interaction effects with all other variants [39]. Here, each genotypic vector is assumed to have been centered and scaled to have mean 0 and standard deviation 1, respectively. Also note that while we limit ourselves to the task of identifying second order (i.e. pairwise) epistatic relationships between genetic variants (i.e. **x**_*r*_ ○ **x**_*s*_), extensions to higher-order epistatic effects are straightforward to implement [39, 60].

There are a few assumptions that need to be made in order to ensure model identifiability in Equation (2). First, because we consider application scenarios where *p > n*, we recall our previous approach and assume that individual effect sizes follow univariate normal distributions, or *β_s_ ~ N* (0, *ω*^2^*/*(*p −* 1)) and *θ_s_ ~ N*(0*, σ*^2^*/*(*p* − 1)) for *s ≠ r* [39]. With the assumption of normally distributed effect sizes, the model defined in Equation (2) is equivalent to a variance component model where **m**_*r*_ ~ N(**0**, *ω*^2^**K**_*r*_) with 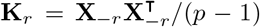 being the genetic relatedness matrix computed using genotypes from all variants other than the *r*-th; and **g**_*r*_ ~ N (**0**, *σ*^2^ **G**_*r*_) with **G**_*r*_ = **D**_*r*_**K**_*r*_**D**_*r*_ representing a relatedness matrix computed based on pairwise interaction terms between the *r*-th variant and all other variants. Here, **D**_*r*_ = diag(**x**_*r*_) denotes an *n × n* diagonal matrix with the genotype vector of interest as its diagonal elements. It is important to note that both **K**_*r*_ and **G**_*r*_ change with every new marker *r* that is considered. For the second assumption, the mean and variance of the liability **l** determine a known disease prevalence *γ* in the population — however, these two parameters are not identifiable from each other given only *γ*. As a result, we follow previous approaches by constraining the variance of **l** to be one (i.e. *ω*^2^ + *σ*^2^ + *τ* ^2^ = 1) [57].

We will formally refer to the liability threshold mixed model defined in Equations (1) and (2) as LT-MAPIT. This approach properly handles the binary nature of case-control data through the direct modeling of **y**. Furthermore, it also appropriately accounts for case-control ascertainment through the direct modeling of liability **l**. The difference between the proportion of cases observed in the population versus those sampled in the case-control study, is accounted for by the distributions of the latent liability values in the population versus in the study. Furthermore, the variance component *σ*^2^ represents a measure on the marginal epistatic effect for the *r*-th variant. Therefore, testing the null hypothesis *H*_0_ : *σ*^2^ = 0 will allow us to examine whether the *r*-th SNP interacts with any other SNPs to exhibit a significant pairwise epistatic interaction.

### Liability Threshold Mixed Model Approximations

In order to estimate and test *σ*^2^ in LT-MAPIT, we first write down the full likelihood of the liability threshold mixed model as a joint function of the variance components (*ω*^2^, *σ*^2^, *τ* ^2^)

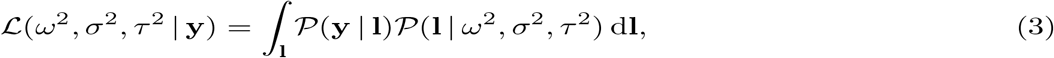

where *P*(**y | l**) is the conditional probability of the observed case-control status **y** given the latent liability **l** as defined in Equation (1), and *P*(**l |** *ω*^2^, *σ*^2^, *τ*^2^) denotes the probability of the latent liability given all other parameters as defined in Equation (2). The likelihood function in Equation (3), unfortunately, involves an *n*-dimensional integration over **l** that cannot be solved analytically. Consequently, we have to rely on approximation methods for inference. Here, we follow previous works [62–65] and approximate the likelihood function with a first order Taylor expansion equated at an estimate of the posterior mean for the liabilities, say **l̂** = E[**l |y**]. Note that the posterior distribution of the liabilities is given as (**l | y**) ∝ (**y | l**) P (**l** *ω*^2^, *σ*^2^, *τ*^2^), which is proportional to the integrand in Equation (3). The resulting approximate likelihood is then written as

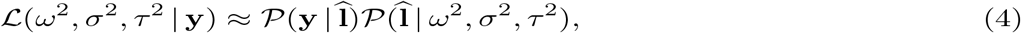

where the first term *P* (**y | l̂**) can be ignored as the posterior mean **l̂** satisfies the constraint set by **y**. The above approximated likelihood may be interpreted as the likelihood of a linear mixed model with the posterior mean estimate **l̂** as the response variable. Therefore, the liability threshold mixed model defined in Equations (1) and (2) can be effectively approximated by a linear mixed effects model on **l̂**, where

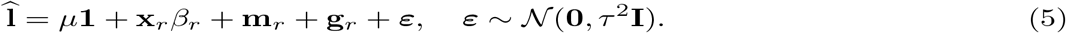

In practice, the posterior mean estimates of the liabilities can be obtained by using a Gibbs sampler, which is an iterative algorithm that generates posterior samples from conditional distributions [66–68]. Thus, given the current parameter estimates of (*ω*^2^, *σ*^2^, *τ*^2^), new values of **l̂** are generated by taking draws from the posterior distribution *P*(**l | y**). These generated values of **l̂** can then be used further to refine the estimates of (*ω*^2^, *σ*^2^, *τ*^2^) through optimizing the approximated likelihood defined in Equation (5). Hence, parameter estimates of (*ω*^2^, *σ*^2^, *τ*^2^) can also be obtained through an iterative optimization procedure. As previously mentioned [63], optimizing a multivariate truncated normal distribution P(**l | y**) to obtain **l̂** is, unfortunately, not straightforward. As a result, instead of a formal Gibbs sampler, we apply a second approximation based on the properties of GWAS data to obtain **l̂** [65]. Specifically, we assume that individuals are essentially unrelated in the sense that **K ≈ I**. We also assume that both the additive and the marginal epistatic effects are small (which is typical for most complex traits [60]), such that that we can ignore the terms **x**_*r*_β_*r*_, **m**_*r*_, and **g**_*r*_. With these additional assumptions, the multivariate normal distribution *P*(**l** *| ω*^2^, *σ*^2^, *τ* ^2^) defined in Equation (2) is approximated by a joint product of independent normal distributions *l_i_* ~ *N*(*μ*, 1). We then apply Markov Chain Monte Carlo (MCMC) to obtain the posterior estimates of **l**. Specifically, in each iteration *t*, posterior samples of **l** are generated from constrained conditional univariate normal distributions

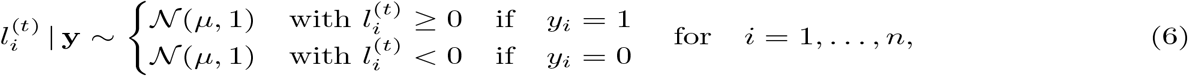

from which **l̂** is analytically computed by averaging over samples drawn during the MCMC iterations. The parameter *μ* = Φ^*−*1^(*γ*) represents the truncation (liability) threshold determined by the known disease prevalence *γ* in the population, with Φ^*−*1^(•) being the inverse cumulative probability density of the standard normal distribution. For all examples in the text, we follow previous approaches [59] and take **l̂** to be a single instance drawn from the posterior in Equation (6).

### Parameter Estimation and Variance Component Test

With the approximations described in the previous section, we are now at a stage to perform joint estimation of the all variance component parameters in the LT-MAPIT model. To do so, we make use of the MQS approach for parameter estimation and hypothesis testing [60]. Briefly, MQS is based on the computationally efficient method of moments and produces estimates that are mathematically identical to the Haseman-Elston (HE) cross-product regression [69]. To estimate the variance components with MQS, we first multiply Equation (5) by a variant specific projection (hat) matrix **H**_*r*_ onto both the null space of the intercept and the corresponding genotypic vector **x**_*r*_. Namely, 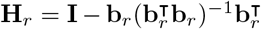 with **b**_*r*_ = [**1**, **x**_*r*_]. After this projection procedure, we obtain a simplified liability threshold mixed model

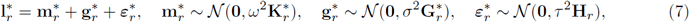

where 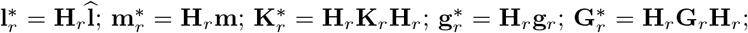 and 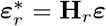, respectively. Then for each variant considered, the MQS estimate for the marginal epistatic effect is computed as

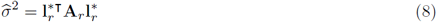

where 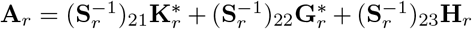 with elements 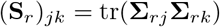 for the covariance matrices subscripted as 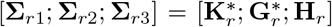. Here, tr(•) is used to denote the matrix trace function. It has been shown that a given marginal epistatic variance component estimate *σ̂*^2^ follows a mixture of chi-square distributions under the null hypothesis [39]. Namely, 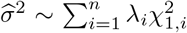, where 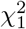 are chi-square random variables with one degree of freedom and (λ_1_, … |, λ_n_) are the eigenvalues of the matrix

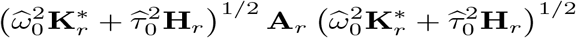

with 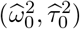 being the MQS estimates of (*ω*^2^, *τ*^2^) under the null hypothesis. Several approximate and exact methods have been suggested to obtain p-values under the distribution of *σ̂*^2^. One frequented choice is Davies exact method [70, 71], which we will use here.

Because of the approximations we use when specifying LT-MAPIT (in the presence of case-control ascertainment), the MQS based testing procedure can result in conservative p-values — even when the model fit is conducted on the liability scale [72]. To circumvent the conservative p-value issue, we propose a recalibration approach following the main idea of genomic control [73] by using the following readjusted test statistics. Specifically, we denote **p̃** = (*p̃*_1_, …, *p̃_p_*) as a vector of p-values computed by using Davies exact method. We consider the genomic control transformation

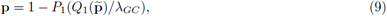

where *Q*_1_(•) denotes the quantile function of the standard chi-square distribution with one degree of freedom; *P*_1_(•) denotes the cumulative density function of the standard chi-square distribution with one degree of freedom; and *λ_GC_* = *m̂ /m̃* is a genomic control factor with *m̂* being the median of *Q*_1_(**p***̃*) and *m̃* ≈ 0.455 being the median of the standard chi-square distribution [74–76]. With this readjustment, the ranking of variants that are most likely to be involved in epistatic interactions remains the same. However, the readjusted p-values now become well-calibrated, which we demonstrate later via simulations. All MAPIT-derived results that we present throughout the rest of the paper will be based on the readjusted marginal epistatic statistics.

### Software Availability

Code for implementing LT-MAPIT is freely available within the “MArginal ePIstasis Test” (MAPIT) software located at https://github.com/lorinanthony/MAPIT. All MAPIT functions use the Com-pQuadForm R package to compute p-values with the Davies method. Note that the Davies method can sometimes yield a p-value that exactly equals 0. This can occur when the true p-value is extremely small [77]. In this case, we report p-values as approximately 0. If this is of concern, practitioners can compute p-values for all MAPIT based functions using Kuonen’s saddlepoint method [77, 78] or Satterthwaite’s approximation equation [79].

### Competing Methods

In the simulations and real data applications presented in this paper, we compare the LT-MAPIT model to five different epistatic mapping methods. The first is the original MAPIT method [39], which treats the case-control labels as a quantitative trait by fitting a linear mixed model directly onto the binary response variable (following the argument that the first order Taylor series expansion is an approximation to a generalized linear model [80–82]). The second method we consider reframes the MAPIT framework within the context of a generalized linear mixed model (GLMM) with a canonical logit link function (MAPIT-Logit). Briefly, the MAPIT-Logit takes an alternative route for GLMM inference and identifies marginal epistatic effects using a penalized quasi-likelihood (PQL) approach (specific details in Supporting Information) [83–85]. The last three methods we use for model comparisons are variations of the PLINK epistatic mapping software (version 1.9) [86]. Here, we use the --`epistasis` argument to fit an exhaustive search (generalized) linear model, which iteratively tests all possible pairwise interactions to identify the exact pairs of variants involved in significant epistasis [48, 49]. Specifically, we consider the following specifications for PLINK: (a) the standard linear regression model where the binary class labels are again treated as quantitative traits, (b) the standard logistic regression model for case-control phenotypes (PLINK-Logit), and (c) a liability threshold exhaustive search model (LT-PLINK). In the last case, we simply fit the linear regression version of PLINK with the same estimated liability values that are given to LT-MAPIT. The PLINK software is publicly available at https://www.cog-genomics.org/plink2/epistasis.

### Simulated Data Generation

To validate LT-MAPIT in terms of its ability to preserve type I error control, and to assess its power compared to the aforementioned competing methods, we carried out a series of simulation studies under a wide range of configurations. Each of these experiments make use of synthetic genotypes that are independently generated to have *p* = 6000 SNPs with allele frequencies randomly sampled from a uniform distribution over values ranging from [0.05, 0.5]. We do not simulate these variants in linkage disequilibrium, partly because of the heavy computational burden involved in generating correlated genotypes under the liability threshold model setting [58], and partly because it has been shown that the distribution of genetic similarity matrices is not affected much by its presence [87]. These genotypes are simulated for a population of size *n* = 2.5 *×* 10^6^. Phenotypes are then generated on the continuous latent liability scale. Disease status for each individual is determined by comparing each simulated *l_i_* with respect to a threshold of liability that is predetermined by the population prevalence parameter *γ*. In this work, we consider three levels of prevalence *γ* = 0.1%, 0.25%, and 0.5%. For each simulated run, an equal number of (randomly selected) cases and controls contribute to the case-control sample. All simulations are replicated 100 times and used a total of *n* = 5000 individuals (2500 cases and 2500 controls).

### Preprocessing of WTCCC GWAS Data

The Wellcome Trust Case Control Consortium (WTCCC) 1 study GWAS data [61] (http://www.wtccc.org.uk/) consists of about 14,000 cases of seven common diseases, including 1,868 cases of bipolar disorder (BD), 1,926 cases of coronary artery disease (CAD), 1,748 cases of Crohn’s disease (CD), 1,952 cases of hypertension (HT), 1,860 cases rheumatoid arthritis (RA), 1,963 cases of type 1 diabetes (T1D) and 1,924 cases of type 2 diabetes (T2D), as well as 2,938 shared controls. We selected an initial set of 458,868 total shared SNPs following the quality control procedures of previous studies [28, 82, 88]. Then for each trait, we retained final data sets with about 363,000 SNPs that had a minor allele frequency (MAF) above 0.05. The disease prevalence *γ* for each trait was determined by referencing past work [89–95] (see Table S1 in Supporting Information).

In the WTCCC GWAS data set, we perform three sets of analyses. The first involves using LT-MAPIT with the usual genome-wide genetic relatedness matrix, where for each SNP *r* in turn, the matrix 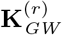 is computed using all other SNPs *s* ≠ *r* in the study. The second analysis involves using LT-MAPIT with a *cis*-genetic relatedness matrix, where alternatively for each SNP *r*, the matrix 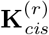 is computed using only variants within a 1 Mb window of SNP *r* on the same chromosome. The last analysis is centered around fitting LT-MAPIT with a genetic relatedness matrix 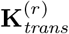, which is computed using only the corresponding *trans*-SNPs located outside of the previously defined 1 Mb *cis*-window for a given SNP *r*. In each of the analyses, SNPs are mapped to the closest neighboring gene(s) using the dbSNP database provided by the National Center for Biotechnology Information (http://www.ncbi.nlm.nih.gov/SNP/).

## Results

### Null Simulations and Type I Error Control

We make use of the following simulation scheme in order to investigate whether LT-MAPIT preserves the desired type I error rate and produces well-calibrated p-values under the null hypothesis. In these numerical experiments, we begin by simulating *p* = 6000 SNPs for a population of *n* = 2.5 × 10^6^ individuals, and generate their liability phenotypes using a linear regression model based on various genetic architectures. Specifically, with these simulated genotypes, we randomly select 1000 causal additive SNPs and draw their corresponding effect sizes from a standard normal distribution. Next, we create residual errors also from a standard normal distribution. Both the genetic effects of the causal SNPs and the effects of the residual errors are scaled to ensure a narrow-sense heritability of 60% on the liability scale. Afterwards, based on predetermined population prevalence levels *γ* = {0.1%, 0.25%, 0.5%}, we select the *γ* proportion of individuals who have the largest liability values to be cases and the remaining to be controls. We then randomly select 2500 cases and 2500 controls as our case-control data to perform analysis. For each *γ*, we performed 500 simulation replicates. Note that, due to the desired experimental property of case-control ascertainment (and the consequent necessary large population size), we are unable to use real genotypes and hence make use of synthetic data.

In these simulations, the idea of the null model holds because there are no interaction effects present during the generation of phenotypes, and LT-MAPIT solely searches for significant marginal epistatic effects that are a summation of pairwise interactions. All evaluations of test calibration and type 1 error control are strictly based on the liability threshold mixed model fitting algorithm presented in the Equations (5)-(8); and the p-values used for model assessment correspond to the readjustment procedure proposed in Equation (9). Figure 1 shows the quantile-quantile (QQ) plots based on the distribution under the null hypothesis. Similarly, Table 1 shows the empirical type I error rates estimated for LT-MAPIT at significance levels *α* = 0.05, 0.01, 0.001, and 0.0001, respectively. Despite being slightly liberal in the assignment of p-values at smaller prevalences (i.e. *γ* = 0.1% case) and at larger critical values (i.e. *α* = 0.05 or 0.01 cases), LT-MAPIT is able to appropriately control the type I error rate for smaller critical values across various prevalences.

**Figure 1.**
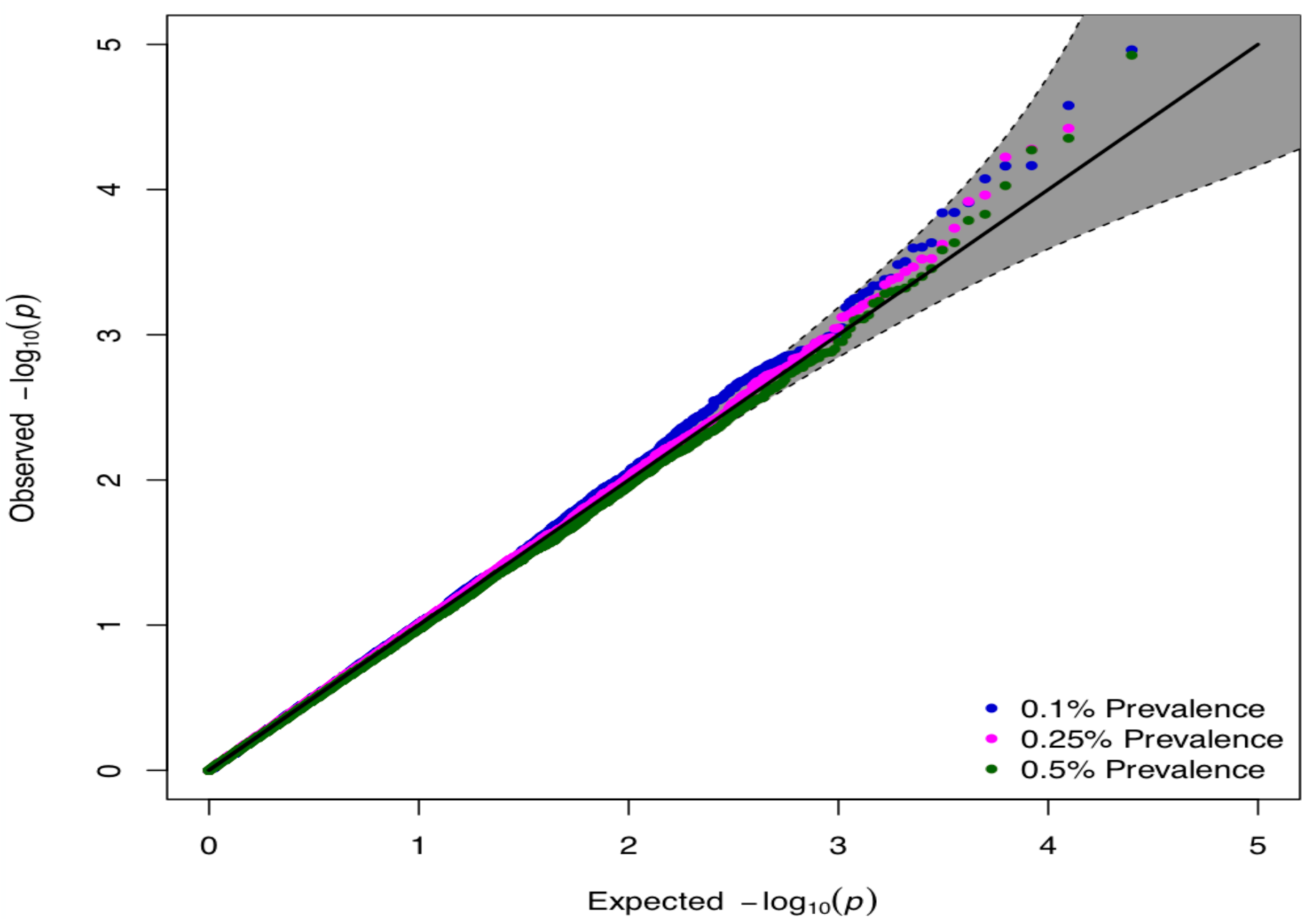
Calibration of p-values produced by LT-MAPIT via QQ-plots. The QQ-plots applying MAPIT to 500 simulated null datasets assuming disease prevalence in the population set to *γ* = 0.1% (blue), 0.25% (pink), and 0.5% (green). The 95% confidence interval for the null hypothesis of no association is shown in grey. Sample size was 5000 (2500 cases, 2500 controls) for all cases.

**Table 1.** Empirical type I error estimates of LT-MAPIT. Each entry represents type I error rate estimates as the proportion of significant p-values under the null hypothesis. Phenotypes were generated on the liability scale with disease prevalence in the population set to *γ* = 0.1%, 0.25%, and 0.5%. Empirical size for the analyses used significance thresholds of *α* = 0.05, 0.01, and 0.001. Sample size was 5000 (2500 cases, 2500 controls) for all cases. Results are based on 500 simulated data sets. Values in the parentheses are the standard deviations across replicates.

| Prevalence ($\gamma$) | $\alpha = 0.05$ | $\alpha = 0.01$ | $\alpha = 0.001$ | $\alpha = 0.0001$ |
| --- | --- | --- | --- | --- |
| 0.1% | 0.0572 (0.0102) | 0.0136 (0.0033) | 0.0010 (0.0012) | 0.0001 (0.0001) |
| 0.25% | 0.0484 (0.0025) | 0.0114 (0.0018) | 0.0008 (0.0013) | 0.0002 (0.0005) |
| 0.5% | 0.0454 (0.0050) | 0.0106 (0.0029) | 0.0008 (0.0013) | 0.0002 (0.0005) |

### Power Assessment and Method Comparisons

The performance of LT-MAPIT approach is evaluated through comparative studies with existing works (see Material and Methods for details). Our goal here is two-fold. First, we want to detect individual variants that are involved in epistasis. Second, we want to explicitly identify pairwise (or higher-order) epistatic interactions. Among many recently proposed epistatic mapping procedures, we focus on comparing LT-MAPIT against two marginal epistasis detection methods (MAPIT and MAPIT-Logit [39]), and three exhaustive search models (PLINK, PLINK-Logit, and LT-PLINK [86]).

We utilize the following simulation design to assess the utility of LT-MAPIT for case-control studies. Here, we again use simulated genotypes to generate continuous phenotypes (on the liability scale) that this time mirror genetic architectures affected by a combination of additive and pairwise epistatic effects. Specifically, we randomly choose 1,000 causal SNPs to directly affect the liability and which classify into three groups: (1) a small set of interaction SNPs, (2) a larger set of interaction SNPs, and (3) a large set of additive SNPs. Here, SNPs interact between sets, so that SNPs in the first group interact with SNPs in the second group, but do not interact with variants in their own group (the same rule applies to the second group). All causal SNPs in both the first and second groups have additive effects and are involved in pairwise interactions, while causal SNPs in the third set only have additive effects. Both the additive and epistatic interaction effect sizes are drawn from standard normal distributions. We scale these genetic effects so that collectively they explain a fixed proportion of the total variance on the liability scale, which again is set to be 60%. In addition, this scaling ensures that the additive effects make up a proportion *ρ* of the total genetic variance, while the pairwise interactions make up the remaining proportion (1 – *ρ*). Altogether, we consider simulations that depend on the three aforementioned parameters:

- *γ*: the assumed disease prevalence in the population, as previously defined;
- (1 – *ρ*): the proportion of genetic variance (on the liability scale) that is contributed by the interaction effects between the first and second groups of causal SNPs;
- *p*_1_*/p*_2_*/p*_3_: the number of causal SNPs in each of the three groups, respectively.

We set *ρ* = {0.5, 0.8}, and define four scenarios corresponding to *p*_1_*/p*_2_*/p*_3_ = 10/10/980 (scenario I), 10/20/970 (scenario II), 10/50/940 (scenario III), and 10/100/890 (scenario IV). In the case where *ρ* = 0.5, the additive and epistatic effects are assumed to equally contribute to the liability phenotypes. In the alternative case, where *ρ* = 0.8, the liabilities are dominated by additive effects. We analyze 100 different simulated datasets for each value of *ρ*, across each of the four scenarios. Unless stated otherwise, all of the results described in the main text are based on the cases in which case-control data are simulated with disease prevalence *γ* = 0.25%. The results for *γ* = 0.1% and 0.5% can be found in Supporting Information (see Figures S1-S4).

Figure 2 shows the empirical power of LT-MAPIT to detect both group 1 and 2 causal variants, respectively, under a Bonferroni corrected genome-wide significance threshold. Overall, we see that LT-MAPIT’s ability to detect epistatic markers mostly depends on the variance explained by the epistatic SNPs. For example, in Figure 2(a), each causal variant in group 1 is expected to explain (1 − *ρ*)*/p*_1_ = 5% of the genetic variance on the liability scale — while in Figure 2(b), these same variants are expected to only explain 2%. In these situations, due to these large effects, LT-MAPIT’s power is greater for detecting group 1 variants in all four scenarios. It is important to note that, unlike in the case for purely quantitative traits (without ascertainment) [39], the liability threshold specification of MAPIT demonstrates an empirical power that also depends on the total interaction effect size. More specifically, LT-MAPIT’s ability to detect group 1 causal epistatic SNPs is slightly affected by the number of group 2 causal SNPs, which also determines the pairwise interaction effect size in simulations. Therefore, LT-MAPIT displays great power when there are only a small number of interactions, each with a large effect size (e.g. scenarios I and II); however, its power marginally decreases in the more polygenic setting where there are a large number of interactions, each with a small effect size (e.g. scenarios III and IV). We hypothesize that this behavior occurs for case-control data because the posterior sampling of the latent liability phenotypes is directly impacted by the magnitude of the pairwise interaction effect size.

**Figure 2.**
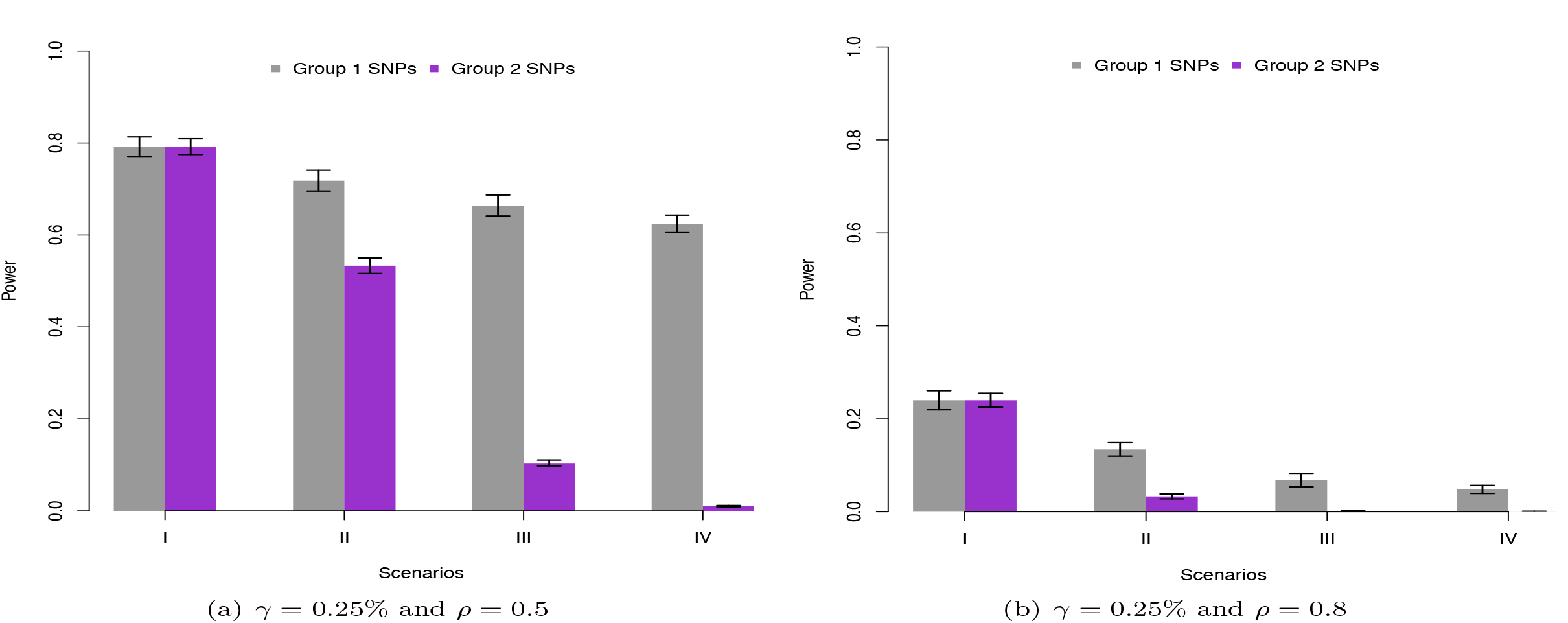
Empirical power to detect simulated causal interacting makers at a genome-wide significance level. Group 1 and 2 causal markers are colored in grey and purple, respectively. Figures (a) and (b) show the power of LT-MAPIT to identify SNPs in each causal group under the Bonferroni-corrected significance level *α* = 8.3 × 10^*−*6^. Phenotypes were generated on the liability scale with disease prevalence in the population set to *γ* = 0.25%. Here, the parameters *ρ* = 0.5 in Figure (a) and *ρ* = 0.8 in Figure (b) are used to determine the proportion of phenotypic variance (on the liability scale) that is contributed by interaction effects. Results are based on 100 simulated replicates and the lines on each bar represent a 95% standard error interval due to resampling.

Figure 3 depicts the direct comparison between the MAPIT-based approaches and the PLINK-based exhaustive searches to accurately identify marginal epistatic markers in each of the two causal groups. Here, we assess power by comparing each model’s true positive rate (TPR) at a 1% false positive rate (FPR). We consider the marginal epistatic detection in the PLINK exhaustive searches by first ordering the resulting p-values for all possible interactions, and then computing the power for correctly identifying unique causal SNPs. For example, if the top p-values from an exhaustive search include interactions SNP1-SNP2, SNP2-SNP3, SNP4-SNP5, and only SNP2 is the true causal epistatic variant, then the top three pairs marginally identify only one true variant and four false variants. Under this evaluation, the methods that do not account for ascertainment (MAPIT, MAPIT-Logit, PLINK, and PLINK-Logit) consistently perform the worst. LT-MAPIT constantly performs the best when the proportions of phenotypic variance explained by additive and epistatic effects are equal (i.e. *ρ* = 0.5). However, when additivity dominates this variance, LT-MAPIT’s advantage over LT-PLINK depends on the disease prevalence. When *γ* is small, LT-MAPIT maintains comparable power; while, LT-PLINK gains an advantage as *γ* increases.

**Figure 3.**
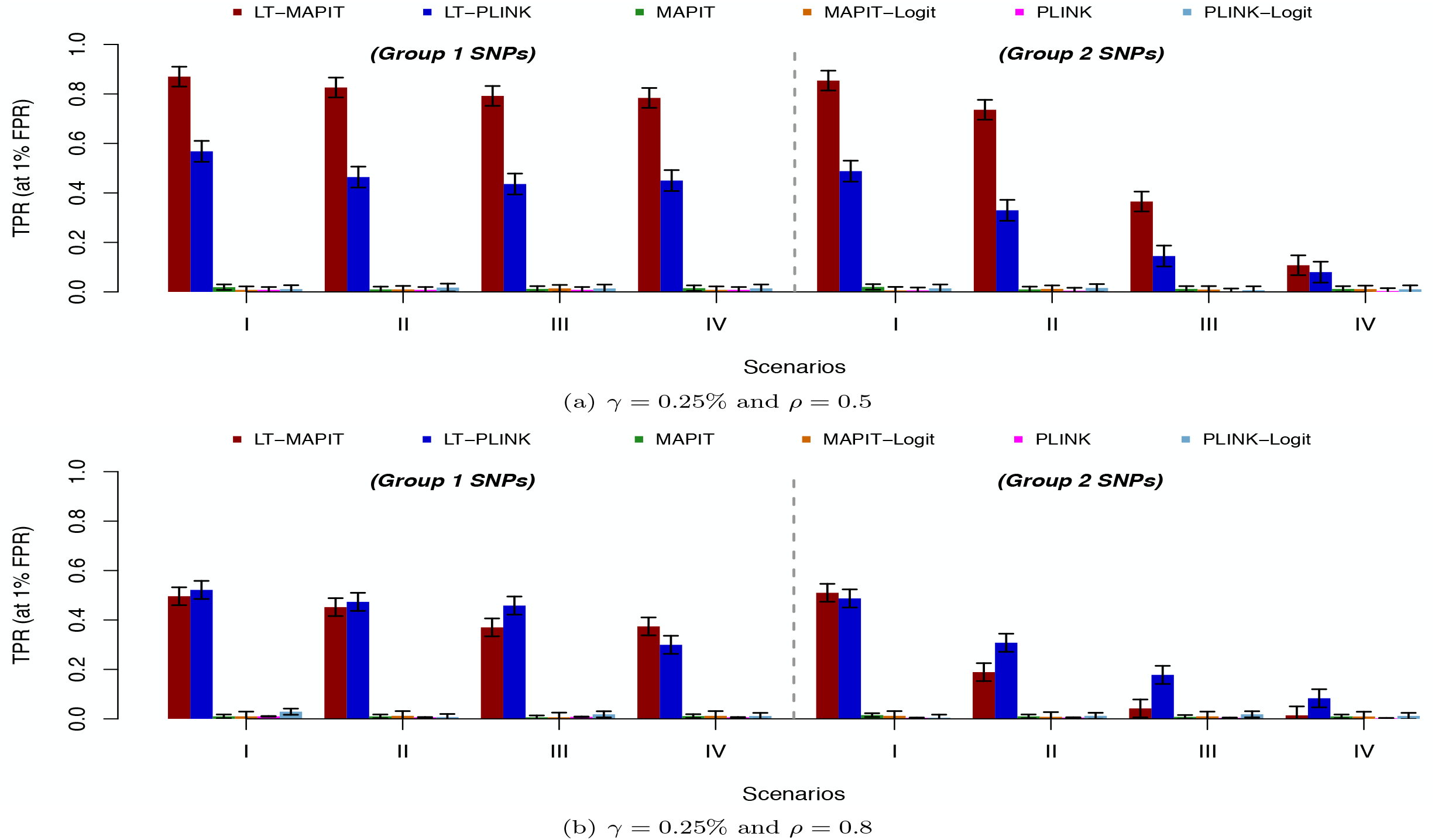
Power comparison for detecting group 1 and group 2 causal SNPs. The association mapping ability of LT-MAPIT (red) is compared to five competing methods: (i) the original linear mixed model version of MAPIT (green), (ii) the binary mixed model MAPIT-Logit (orange) fit by using penalized quasi-likelihoods (PQL), (iii) the linear exhaustive search model PLINK (pink), (iv) the generalized linear model PLINK-Logit (light blue), and (v) the liability threshold model LT-PLINK (blue). Each approach is assessed under four simulation scenarios. Phenotypes were generated on the liability scale with disease prevalence in the population set to *γ* = 0.25%. Here, the parameters *ρ* = 0.5 in Figure (a) and *ρ* = 0.8 in Figure (b) are used to determine the proportion of phenotypic variance (on the liability scale) that is contributed by interaction effects. The y-axis gives the rate at which true causal variants were identified (TPR) at a 1% false positive rate (FPR). Results are based on 100 simulated replicates and the lines on each bar represent a 95% standard error interval due to resampling.

In addition to detecting marginal epistatic effects, we also assessed each method’s ability to explicitly identify significant interactions. Specifically, we investigate the use of LT-MAPIT as the initial step in a two-step prioritization based association mapping procedure [39]. Under this type of approach [45,49,51], we first apply LT-MAPIT as a single-SNP test to identify associated genetic variants with non-zero marginal epistatic effects, and then focus on the significantly identified variants to test all pairwise interactions between them. Figure 4 compares the utility of this prioritization procedure when selecting the top *υ* = {50, 100, 250, 500} SNPs as opposed to ranking significant interactions estimated via the liability threshold exhaustive search LT-PLINK. Altogether, just searching over the top 250 prioritized SNPs using LT-MAPIT provides more power in finding true pairwise epistatic interactions across all scenarios. This holds true even for most of the cases in which the additive effects contribute to a majority of the variance on the liability scale.

**Figure 4.**
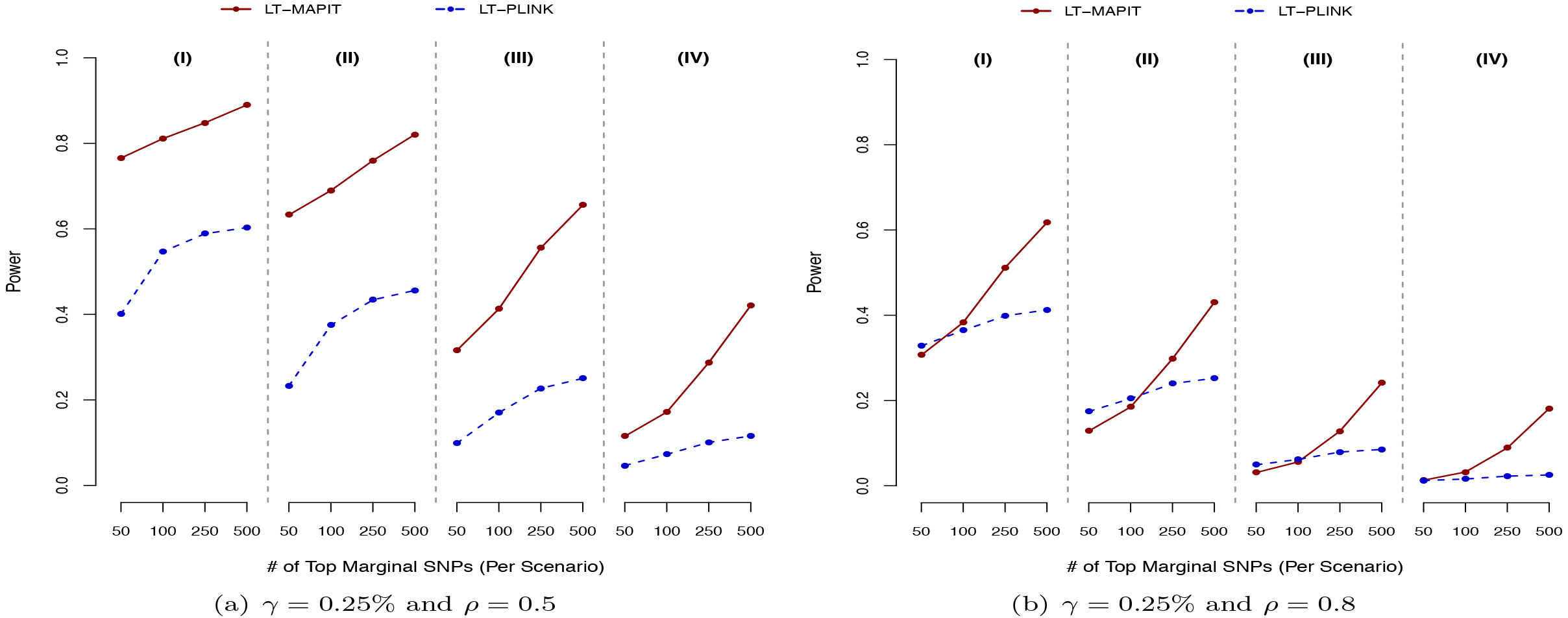
Power comparison with exhaustive search procedures to detect epistatic pairs. LT-MAPIT (red) is assessed as an initial step in a pairwise epistatic prioritization process compared against the more conventional exhaustive search LT-PLINK (blue), which serves as a baseline. While using LT-MAPIT, the search for epistatic pairs occurs between the top ranked {50, 100, 250, 500} significant marginally associated SNPs. Both methods are evaluated under four simulation scenarios. Phenotypes were generated on the liability scale with disease prevalence in the population set to *γ* = 0.25%.

Lastly, we check method sensitivity to correct prevalence specification. To evaluate this, we run a final simulation study where we set the true disease prevalence in the population to be *γ* = 0.25%, but then generate latent liabilities for LT-MAPIT and LT-PLINK using misspecified thresholds based on prevalences *γ̃* = 0.1% (low) and *γ̃* = 0.5% (high). In Supporting Information (see Figures S5-S6), we show that LT-MAPIT is generally more robust to a liberal misspecification of the ascertainment threshold (i.e. *γ̃* = 0.5%), as opposed to a conservative one (i.e. *γ̃* = 0.1%). This is particularly apparent in the *ρ* = 0.8 simulation cases where additive effects dominant the phenotypic variance. While the performance of LT-PLINK is seemingly unaffected by the misspecification of the liability threshold when it comes to identifying marginal epistatic variants, it also follows the same liberal versus conservative behavior as LT-MAPIT when detecting explicit epistatic pairs (see Figure S6). Nonetheless, LT-PLINK still underperforms overall when compared to LT-MAPIT in most of the simulation scenarios we consider.

### Practical Application to WTCCC Data

In addition to our simulation study, we further assess LT-MAPIT’s ability to detect marginal and pairwise epistasis in the seven complex diseases provided by the Wellcome Trust Case Control Consortium (WTCCC) 1 study [61]. Briefly, these diseases include: bipolar disorder (BD), coronary artery disease (CAD), Crohn’s disease (CD), hypertension (HT), rheumatoid arthritis (RA), type 1 diabetes (T1D), and type 2 diabetes (T2D) — where the disease prevalence *γ* for each trait has been previously documented [89–95]. For an in-depth analysis on the summed epistatic effects involving any given SNP, we implement LT-MAPIT using three distinct genetic relatedness matrices: (i) **K**_*GW*_ which tests against all SNPs genome-wide, (ii) **K**_*cis*_ which tests against all *cis*-SNPs within a given 1 Mb genomic window, and (iii) **K**_*trans*_ which tests against all other SNPs located outside the *cis*-window (details in Materials and Methods). Intuitively, using **K**_*cis*_ will be more powerful than using either **K**_*trans*_ or **K**_*GW*_ if epistatic interactions are more likely to happen between *cis*-SNP pairs, rather than between SNPs that are located genome-wide — however, less powerful otherwise.

For each trait, we provide summary tables which list the marginal epistatic p-values for all SNPs when the LT-MAPIT model is fit using the different genetic relatedness matrices (see Table S2-S8 in Supporting Information). Corresponding QQ-plots of the– log_10_(*P*) transformed p-values visually display this information in Figure 5. To contrast LT-MAPIT’s marginal association findings, we again directly compare results from the liability threshold exhaustive search model LT-PLINK. The first half of Table S9 details comparisons between these two approaches and their effectiveness in finding marginally epistatic variants across all seven traits. We are reminded that LT-MAPIT searches over a reduced space, and thus only requires *p* tests for a data set with *p* genetic markers. Therefore, a significant marginal association for a particular SNP identified by LT-MAPIT is determined by using a (traditional) trait specific Bonferroni-corrected significance p-value threshold *P* = 0.05*/p_d_*, where *p_d_* is the number of SNPs for disease trait *d*. For LT-PLINK, a single SNP was deemed marginally significant if it belonged to an epistatic pair with a p-value below the genome-wide threshold *P* = 0.05*/C*(*p_d_*, 2) — where *C*(•, 2) denotes the binomial coefficient enumerating all possible SNP pairs. Note that this is equivalent to implementing a Bonferroni-corrected threshold for exhaustive search methods [49].

**Figure 5.**
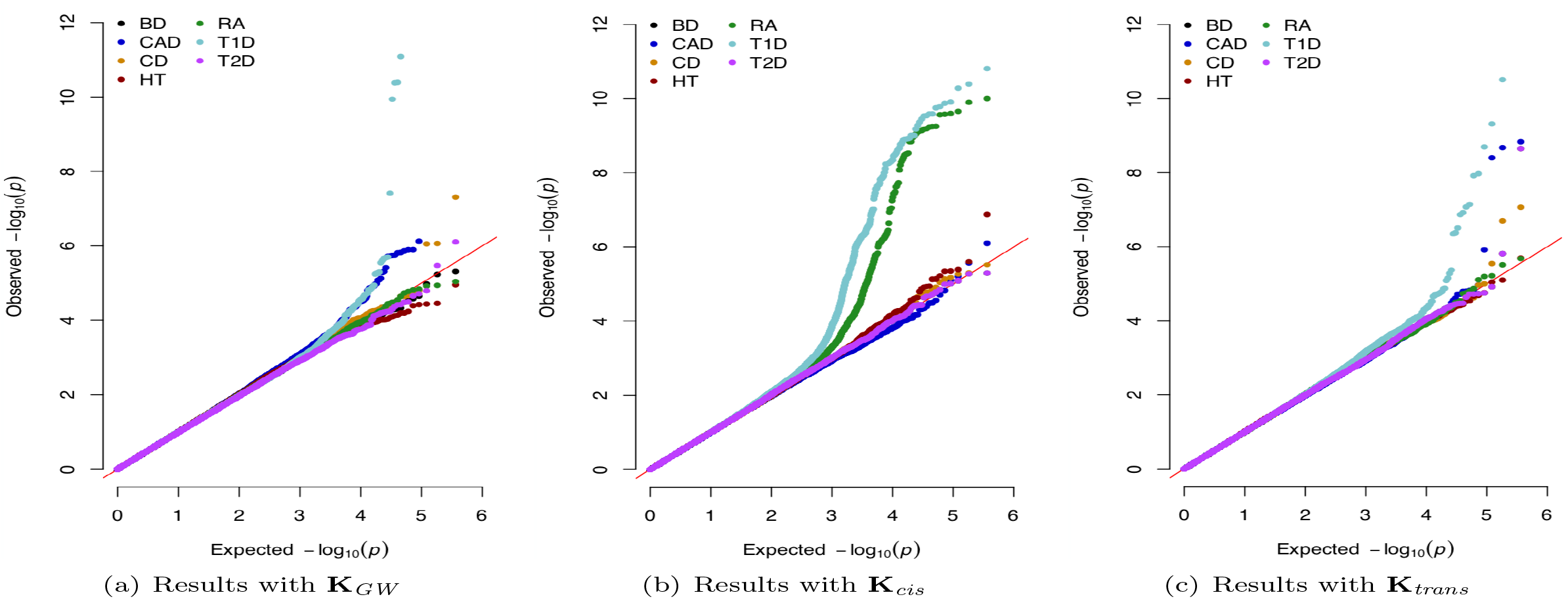
QQ-plots of LT-MAPIT p-values in the WTCCC data set. The seven diseases analyzed include: bipolar disorder (BD), coronary artery disease (CAD), Crohn’s disease (CD), hypertension (HT), rheumatoid arthritis (RA), type 1 diabetes (T1D), and type 2 diabetes (T2D). Each figure corresponds to using LT-MAPIT with a different genetic relatedness matrix. Namely, Figure (a) corresponds to fitting the liability threshold mixed model with a genetic relatedness matrix **K**_*GW*_, which is computed using all SNPs in the study. Figure (b) corresponds to using **K**_*cis*_, which is computed using only variants within a 1 Mb window (on the same chromosome) from the SNP of interest. Figure (c) are results from using **K**_*trans*_, which is computed using only the corresponding *trans*-SNPs located outside of the previously defined 1 Mb *cis*-window.

In Table 2, we list all genes found by LT-MAPIT with at least two SNPs having marginal epistatic p-values that satisfy the Bonferroni-corrected significance cutoffs. Overall, LT-MAPIT was able to identify noteworthy SNPs found within the coding regions of genes that are known to have relevant associations. For example, in BD and RA, the genome-wide and *trans*-based analyses detected variants with the lowest marginal epistatic p-values near the gene *CD200*. In the context of BD, *CD200* is known to be involved in microglial activation and plays a pivotal role in pathways connected to psychiatric disorders [96, 97]. Alternatively, when studying RA, irregular expression of *CD200* has been shown to contribute to abnormal Th17 cell differentiation [98], regulate myeloid cell function [99], and has been proposed as a potential therapeutic target [100, 101]. For HT, the *cis*-window version of LT-MAPIT identified significant variants near the gene *PSAT1*, whose low expression levels serve as a biomarker of decreased serine biosynthesis pathway activity and increased risk for pulmonary hypertension [102, 103]. Similarly, *IFNB1* has been validated as a gene that affects immunity in T1D [104–106] and is a key member of the inflammatory signature for progressive diabetic nephropathy in T2D [106, 107].

**Table 2.** Gene coding regions with at least two SNPs having marginal epistatic p-values that satisfy Bonferroni-corrected thresholds in the analysis of the WTCCC data set. The seven diseases analyzed include: bipolar disorder (BD), coronary artery disease (CAD), Crohn’s disease (CD), hypertension (HT), rheumatoid arthritis (RA), type 1 diabetes (T1D), and type 2 diabetes (T2D). Listed for all regions are the SNPs with the lowest p-values. The reference column gives literature sources that have previously suggested some level of association between a given gene and disease. Columns 4-6 correspond to using LT-MAPIT with different genetic relatedness matrices. Namely, these include: the genome-wide matrix **K**_*GW*_ (GW), a *cis*-based matrix **K**_*cis*_ (Cis), and a *trans*-based matrix **K**_*trans*_ (Trans), respectively. : Multiple SNPs in the MHC gene class regions are significant, so a single SNP with the lowest marginal epistatic p-value is reported as the most extreme. There are three subregions that divide the MHC region: the class I region (29.8Mb-31.6Mb), the class III region (31.6Mb-32.3Mb), and the class II region (32.3Mb-33.4Mb).

| Disease | Chromosome | Nearest Gene(s) | P-Value (GW) | P-Value (Cis) | P-Value (Trans) | Reference(s) |
| --- | --- | --- | --- | --- | --- | --- |
| BD | 3 | CD200 | $\approx 0$ (rs6438075) | — | $\approx 0$ (rs6438075) | [96, 97] |
| CAD | 1 | LMX1A | $1.62 \times 10^{-14}$ (rs16841238) | — | $1.47 \times 10^{-9}$ (rs16841238) | — |
| HT | 9 | PSAT1 | — | $1.32 \times 10^{-7}$ (rs944513) | — | [102, 103] |
| RA | 3 | CD200 | $\approx 0$ (rs6438075) | — | $\approx 0$ (rs6438075) | [98–101] |
| RA | 6 | MHC Class I<sup>§</sup> | — | $8.93 \times 10^{-8}$ (rs9266775) | — | [125, 126] |
| RA | 6 | MHC Class III<sup>§</sup> | — | $1.00 \times 10^{-10}$ (rs9268480) | — | [125, 126] |
| RA | 6 | MHC Class II<sup>§</sup> | — | $5.85 \times 10^{-10}$ (rs2857154) | — | [125, 126] |
| RA | 9 | TPD52L3 | $\approx 0$ (rs2290913) | — | $\approx 0$ (rs2290913) | — |
| T1D | 6 | MHC Class I<sup>§</sup> | — | $5.47 \times 10^{-11}$ (rs261948) | $2.03 \times 10^{-9}$ (rs2523485) | [127, 128] |
| T1D | 6 | MHC Class III<sup>§</sup> | $4.87 \times 10^{-23}$ (rs3132959) | $3.73 \times 10^{-10}$ (rs3131294) | $2.82 \times 10^{-22}$ (rs2736177) | [127, 128] |
| T1D | 6 | MHC Class II<sup>§</sup> | $1.82 \times 10^{-17}$ (rs9272723) | $1.29 \times 10^{-10}$ (rs2858308) | $4.85 \times 10^{-10}$ (rs2647015) | [127, 128] |
| T1D | 9 | IFNB1 | $\approx 0$ (rs6475502) | — | $\approx 0$ (rs6475502) | [104–106] |
| T2D | 1 | LMX1A | $\approx 0$ (rs16841238) | — | — | [129, 130] |
| T2D | 9 | IFNB1 | $\approx 0$ (rs6475502) | — | — | [106, 107] |

As expected from previous statistical epistatic methods that have heavily studied this data [49, 55, 56, 108, 109], LT-MAPIT most noticeably identified the major histocompatibility complex (MHC) region on chromosome 6 in RA and T1D as having many genetic variants that are most likely involved in prominent pairwise interactions. This is clearly visible in Figures S7 and S8 which display manhattan plots of our marginal epistatic genome-wide scans for these two traits. In these figures, spikes across chromosomes suggest loci where members involved in epistatic interactions can be found. Moreover, when strictly considering *cis*-relationships between SNPs via the relatedness matrix **K**_*cis*_, LT-MAPIT was able to detect more marginal signal than LT-PLINK in both RA and T1D (again see Table S9). These results are unsurprising since it is well known that the MHC region holds significant clinical relevance in complex traits and diseases with respect to infection, inflammation, autoimmunity, and transplant medicine [110, 111]. Since this region has also been consistently implicated by other epistatic analyses, we focus our search for significant pairwise interactions to these two phenotypic traits (see the second half of Table S9 for results pertaining to the other five traits).

Similar to what was done in the simulation studies, we utilize the proposed LT-MAPIT prioritization approach to detect explicit epistatic pairs in RA and T1D. To evaluate the power of this method, we first rank SNPs according to their marginal epistatic p-values and then search for significantly associated epistatic pairs between the top *υ* SNPs. Interactions between two variants were then called significant if they had a joint p-value below the Bonferroni-corrected threshold *α* = 0.05*/C*(*υ*, 2). Here, we choose two sets of prioritization thresholds: one low-dimensional threshold *υ* = {10, 50, 100, 200} (see Figure S9(a)) and one high-dimensional threshold *υ* = {500, 1000, 2000, 5000} (see Figure S9(b)). In each case, we compared results with significant pairs detected by exhaustive search via LT-PLINK, as a baseline. Naturally, this comparison serves as a way for us to continue our assessment of LT-MAPIT as a viable prioritization approach within the context of real case-control data. Altogether, the exhaustive search method identified 75 significant epistatic pairs in RA and 2753 epistatic pairs in T1D. As demonstrated in the numerical experiments, using LT-MAPIT (particularly with **K**_*cis*_) efficiently improves upon this power.

## Discussion

We have presented LT-MAPIT for identifying variants that are involved in epistasis in ascertained case-control GWASs. For each variant in turn, LT-MAPIT tests for marginal epistatic effects — the combined epistatic effect between the examined variant and all other variants — to identify variants that exhibit non-zero interactions with another variant without the need to select the specific marker combinations that drive the epistatic association. LT-MAPIT extends the previous MAPIT method for quantitative traits [39] to case-control studies and represents an attractive alternative to conventional epistatic mapping methods [47–49, 56]. With both simulations and real data applications, we have illustrated the benefits of LT-MAPIT.

We have primarily focused on using a liability threshold mixed model to account for case-control ascertainment and the binary nature of case-control data. An alternative approach commonly used for modeling case-control studies and performing variance component test is the logistic mixed model. Both the liability threshold mixed model and the logistic mixed model are prospective models that treat the binary label of case-control status as the outcome variable. The liability threshold mixed model directly accounts for the retrospective sampling in case-control studies by explicitly modeling case-control ascertainment. The prospective logistic model without random effects, on the other hand, is also well known to yield valid likelihood based inference of the odds ratio which is equivalent to a corresponding retrospective logistic model under case-control ascertainment [112]. Unfortunately, the equivalence of the likelihood based inference between a prospective logistic model and a retrospective logistic model does not directly imply an equivalence of likelihood based inference between a prospective logistic mixed model and a retrospective logistic mixed model. Indeed, it is not trivial to derive a corresponding retrospective model for a prospective logistic mixed model, or the other way around, due to case-control ascertainment [113,114]. Consequently, it is also not straightforward to treat the multiple genetic variance components in the prospective logistic mixed model as population level parameters — which we can do with a liability threshold mixed model. Because of the lack of a corresponding retrospective model, it has been shown that the prospective logistic mixed model may not be robust to various modeling misspecifications [115]. Indeed, here we also found that the logistic mixed model does not provide sufficient power in the presence of retrospective sampling and case-control ascertainment. Nevertheless, the inference of logistic mixed model has been well established in the statistics literature [116–119], and the model has been recently applied to perform association tests in case-control studies [120] as well as in RNA sequencing and bisulfite sequencing studies [83–85]. Extending and adapting the logistic mixed model for mapping marginal epistasis would be an interesting avenue to explore in the future.

There are several potential extensions of LT-MAPIT. For example, while we have focused on detecting variants that are involved in pairwise interactions with other variants, LT-MAPIT can be easily extended to detect variants that are involved in higher-order interactions. In particular, we can introduce extra random effects terms to represent the combined higher-order interaction effects between the examined variant and all other variants. Under the normality assumption for the interaction effect sizes, the introduced random effects terms would all follow multivariate normal distributions with the covariance matrices determined as a function of the Hadamard product of the additive genetic relatedness matrix, where the power of the Hadamard product depends on the particular higher-order term one is interested in modeling [33,121,122]. Therefore, in the presence of higher-order interactions, we can apply a multiple variance component model with additional random effects to map these epistatic variants. As another example, we have assumed in LT-MAPIT that the interaction effect between the examined variant and every other variant follows a normal distribution. This normality assumption has been widely applied in many areas of genetics [71, 82, 123] and is known to produce practically unbiased estimates of the total effects, regardless of whether or not the underlying effect sizes actually follow a normal distribution [82, 123]. However, the main idea in LT-MAPIT of mapping marginal epistatic effects is not restricted to the normality assumption for the interaction effect sizes. Indeed, we can incorporate sparsity-inducing priors for effect sizes if the proportion of interaction pairs is known to be small *a priori*, or we can use hybrid effect size priors to accommodate a larger variety of distributions [82, 88]. Different interaction effect size assumptions can be advantageous under different genetic architectures and will likely improve the power of LT-MAPIT further.

LT-MAPIT is not without its limitations. Like other marginal epistatic mapping methods, LT-MAPIT is unable to directly identify detailed interaction pairs despite being able to identify SNPs that are involved in epistasis. However, being able to identify SNPs involved in epistasis allows us to come up with an initial (likely) set of variants that are worth further exploration, and thus represents an important first step towards identifying and understanding detailed epistatic associations. Indeed, we have illustrated a two-step *ad hoc* epistatic association mapping procedure, where we first identify individual SNP associations with LT-MAPIT and then we focus on the most significant associations from the first step to further test all of the pairwise interactions among them in order to identify specific epistatic interaction pairs. Unlike the previous prioritization strategies commonly used in this space, which prioritize SNPs based on additive effects, our two-step procedure is unique in the sense that the SNP set identified in our first step contains SNPs that already display strong epistatic effects with other variants. Therefore, our two-step procedure outperforms alternative prioritization strategies in simulations and real data applications. Nonetheless, we caution that the two-step procedure is still *ad hoc* in nature and could miss important epistatic associations. Exploring statistical approaches that can unify the two steps in a joint fashion would be an interesting area for future research. Besides this main limitation, we also note that LT-MAPIT can be computationally expensive. LT-MAPIT requires fitting a different variance component model for every SNP in turn, and fitting variance component models is known to be computationally challenging [81,124]. In this study, we rely on various approximations to the liability threshold model and the recently developed efficient MQS method to enable efficient variance component test. Compared with the standard REML method, MQS is computationally efficient, allows for accurate p-value computation based on the Davies method, and is statistically produces more accurate estimates than the REML when the variance component is small [60] — a property that is particularly relevant here considering the marginal epistatic effect size is often small in real data. With these techniques, LT-MAPIT is applicable to moderately sized genetic mapping studies with thousands of samples and millions of variants. Still, new algorithms are likely needed to scale LT-MAPIT up to datasets that are orders of magnitude larger.

## Acknowledgements

LC would like to acknowledge the support of Institutional Development Award Number P20GM109035 from the National Institute of General Medical Sciences of the National Institutes of Health, which funds COBRE Center for Computational Biology of Human Disease. XZ would like to acknowledge the support of National Institutes of Health (NIH) Grant R01HG009124 and the support of National Science Foundation (NSF) Grant DMS1712933. This study also makes use of data generated by the Wellcome Trust Case Control Consortium (WTCCC). A full list of the investigators who contributed to the generation of the data is available from www.wtccc.org.uk. Funding for the WTCCC project was provided by the Wellcome Trust under award 076113 and 085475. Any opinions, findings, and conclusions or recommendations expressed in this material are those of the author(s) and do not necessarily reflect the views of any of the funders or supporters.

